# Motility-Driven Viscoelastic Control of Tissue Morphology in Presomitic Mesoderm

**DOI:** 10.1101/2025.10.28.684986

**Authors:** Sahil Islam, Mohd. Suhail Rizvi, Anupam Gupta

## Abstract

During development, embryonic tissues experience mechanical stresses ranging from cellular to supracellular length scales. In response, cells generate active forces that drive rearrangements, allowing the tissue to relax accumulated stresses. The nature of these responses depends strongly on the magnitude and duration of the deformation, giving rise to the tissue’s characteristic viscoelastic behavior. Although experiments have characterized tissue rheology in various contexts, simpler theoretical approaches that directly connect cellular activity to emergent rheological behavior are still limited. In this study, we employ a vertex-based model of epithelial tissue incorporating active force fluctuations in cell vertices to represent cell motility. We capture distinct rounding dynamics and motility-dependent timescales by benchmarking against experimental observations such as the bulging of presomitic mesoderm (PSM) explants driven by Fibroblast Growth Factor(FGF) gradients. Stress relaxation tests reveal rapid short-timescale relaxation alongside persistent long-timescale residual stresses that decrease from anterior to posterior (AP) region of the PSM. By applying oscillatory shear, we analyzed the resulting elastic and viscous responses, revealing motility dependence of storage and loss modulus. Finally, we introduce spatially patterned cues applied in a temporally pulsed manner, mimicking dynamic biochemical or mechanical signals during development. Our results show that while higher motility promotes tissue remodeling in response to these cues, this response is constrained by spatial scale; cellular-scale perturbations are relaxed irrespective of motility strength, preventing complete morphological adaptation.

## I. INTRODUCTION

Embryonic tissues exhibit a complex interplay between mechanical forces and active cellular behaviors during development[1]. In the process, tissue undergoes large-scale deformation and remodeling [2]. As a response, they propagate forces and relax stresses through active cellular rearrangements and passive mechanical resistance largely governed by cytoskeletal mechanics and actomyosin contractility [3–6]. The resulting mechanical responses span a wide range of spatial and temporal scales, making tissues highly sensitive to the nature of the applied deformation [7]. Understanding these principles not only aids in interpreting experimental measurements of spatial variations in mechanical properties, but also forms the basis for predicting how tissues might reorganize and evolve under different developmental or perturbative conditions. Tissue behavior on varying space and time scales has been studied extensively for over a decade. Embryonic chicken tissue, such as neural retina, liver, and heart ventricle, has been shown to behave like a viscoelastic material with elastic relaxation on short timescales and a viscous liquid-like response on longer timescales [8]. More recently, studies have quantitatively characterized the mechanical properties of embryonic tissues using techniques such as ferrofluid microdroplet deformation [2, 9], magnetic bead twisting, and micropipette aspiration [10]. In the presomitic mesoderm (PSM), these approaches have revealed distinct spatial variation in mechanical properties along the anterior–posterior (A–P) axis: both stiffness and viscosity increase from the posterior to the anterior, producing a gradual fluid-to-solid transition across the tissue [2, 3]. Importantly, even at a fixed position along this axis, the mechanical response strongly depends on the timescale of deformation [9]. Localized microdroplet measurements show that tissues initially resist deformation elastically, partially recovering shape on short timescales [2]. Still, over longer durations, stresses relax through cellular rearrangements and junctional remodeling, leading to a viscous-like flow [2, 9]. Thus, the PSM exhibits a biphasic mechanical response, elastic at short times and viscous at long times. This intrinsic timescale-dependent behavior is a hallmark of tissue viscoelasticity in the developing embryo.

In addition to the macroscopic gradients in mechanical properties observed in the PSM, cellular behaviors also vary systematically along the anterior–posterior (A–P) axis. Time-lapse microscopy shows that cells in the anterior region undergo weaker random spatial motion than those in the posterior region [11]. This difference in motility mirrors the gradient of fibroblast growth factor (FGF), a key regulator of cell movement [11], which decreases from posterior to anterior along the PSM [11, 12]. Such coupling between mechanical behavior and cellular activity could be a key determinant of morphogenesis, providing a mechanistic basis for predicting how developing tissues will deform and reorganize under different developmental conditions.

In this work, we use a motile cell-based vertex model [13] to explore how the presomitic mesoderm (PSM) responds to mechanical forces. First, we establish that the model captures key experimental observations, including the characteristic bulging and rounding dynamics of PSM explants in vitro. Using this validated framework, we then extract the viscoelastic timescales that govern tissue mechanics along the anterior–posterior axis. These timescales allow us to make analytical predictions for how the PSM deforms under externally imposed, spatially patterned pulsatile forces. Remarkably, the simulations recapitulate these predictions, providing a direct link between cell motility, viscoelasticity, and large-scale tissue morphology.

## II. ACTIVE VERTEX MODEL

We use the vertex model [13–15], a well-established framework for modeling epithelial tissues, to study the emergence of viscoelasticity and its regulation in embryonic PSM. In the vertex model, individual cells are represented as polygons forming a confluent mesh [Figure S1(a)]. The tissue’s mechanical behavior is described in terms of an energy functional that accounts for deviations from target cell geometry and interfacial tensions:

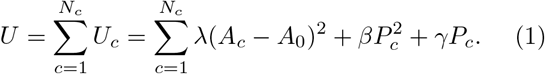

Here, *N*_*c*_ is the total number of cells, *A*_*c*_ and *P*_*c*_ are the instantaneous area and perimeter of cell *c*, and *A*_0_ is the preferred cell area. The parameters *λ, β*, and *γ* respectively control the resistance to area deformations (bulk elasticity), cortical contractility, and effective interfacial tension arising from adhesion and membrane tension. The motion of each vertex follows overdamped dynamics governed by:

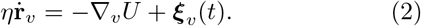

where **r**_*v*_ is the position of vertex *v*, and *η* is an effective friction coefficient that captures viscous resistance from the substrate and/or surrounding fluid. The term ∇_*v*_*U* represents the gradient of the energy *U* taken with respect to vertex position. The stochastic term ***ξ***_*v*_ (*t*) accounts for active movement of cellular junctions and is modeled as Gaussian white noise with zero mean and amplitude proportional to a motility parameter M(**r**) :

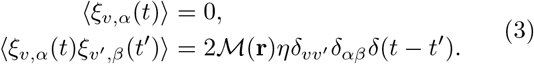

where δ_*vv′*_ ensures that the noise is uncorrelated between different vertices and *ξ*_*v,α*_ is *x* or *y* components of ***ξ***_*v*_ . Together with Eq. 1 and 2, this formulation captures both the mechanical forces that maintain cellular equilibrium and the stochastic, non-equilibrium forces. These stochastic forces effectively represent cellular motility, originating from intracellular processes such as cytoskeletal turnover and actomyosin contractility, and should not be confused with thermal noise [12]. These forces create random movement of cells, which in the context of PSM, parallels Fibroblast Growth Factor (FGF) signaling in coherence with experiments reported in [11, 12] (see Section II for details). Importantly, these random forces at cell vertices drive similar cell–cell contact length fluctuations observed in experiments done on actively fluctuating epithelial junctions [3].

In this model, we also take into account cellular junctional rearrangements, including T1 transitions, where neighboring cells exchange contacts (Figure S1b), and T2 transitions, where cells below a critical size undergo extrusion and are replaced by a multicellular junction (Figure S1c). Cell proliferation is not included in this model, as the simulated timescales are significantly shorter than those of cell division.

## III. ACTIVE VERTEX MODEL REPRODUCES EXPERIMENTAL OBSERVATION INVOLVING PSM EXPLANTS

To benchmark our simulation parameters (*λ, β, γ*) with PSM, we have reproduced an experimental result reported in [16], in which an embryonic PSM cultured ex vivo exhibits spontaneous shape remodeling. Experimental studies have shown that isolated PSM explants spontaneously remodel their shape and fluidize over time, suggesting a tight coupling between motility and mechanical relaxation. By tuning our parameters to capture these behaviors, we establish a consistent framework for exploring how spatial variations in motility and mechanical resistance along the anterior–posterior axis shape tissue morphogenesis.

### A. Cell motility drives bulging of PSM explant

The PSM exhibits a gradient in cellular motility along its anterior-posterior axis, regulated by Fibroblast Growth Factor (FGF) signaling [11, 12]. The highly motile cells in the posterior PSM are able to overcome intrinsic contractile and adhesive forces and adapt a circular shape [Fig. 1(a)]. In contrast, the anterior region, with lower motility, fails to circularize, resulting in a characteristic pear-like tissue shape [Fig. 1(a)]. Notably, this morphological transformation occurs on timescales shorter than those of cell division [16]. To gain deeper insight into the spatial variation of mechanical properties along the anterior-posterior (AP) axis, Arthur et al.[16] isolated tissues from different regions of the presomitic mesoderm (PSM) and allowed them to evolve freely. While both explants ultimately adopted a circular shape, they did so over significantly different timescales [Fig.1(c)]. This rounding up of a tissue is a hallmark of fluidization as fluids try to minimize the surface energy by forming a circular shape. These experiments reveal that motility can drive tissue fluidization, which can control PSM morphogenesis. We have reproduced these experiments using the vertex model where starting from a rectangular tissue, we incorporate FGF gradient by making the strength of motility ( ℳ (**r**)) decay exponentially from posterior to anterior direction with a length scale established by experimental observations [11, 12, 16]. To be consistent with the experiments, we start with an initial tissue state that, due to contractility and the adhesive nature of the cells, shrinks in size initially, but over longer times, motile cells easily overcome the energy barrier required for junctional rearrangement, which facilitates fluidization. This motility-driven fluidization varies along the A-P axis. The posterior PSM being in a higher state of motility becomes rounded in shape quicker than the anterior part, ultimately leading to the emergence of the characteristic pear-like morphology with a decrease in total tissue length[Fig. 1(b)]. We also simulated tissue evolution independently under three different motility strengths. Consistent with experimental findings, we observe that tissues mimicking the anterior region reach circularity more rapidly than those representing the middle or posterior PSM [Fig. 1(d,e)].

**FIG. 1.**
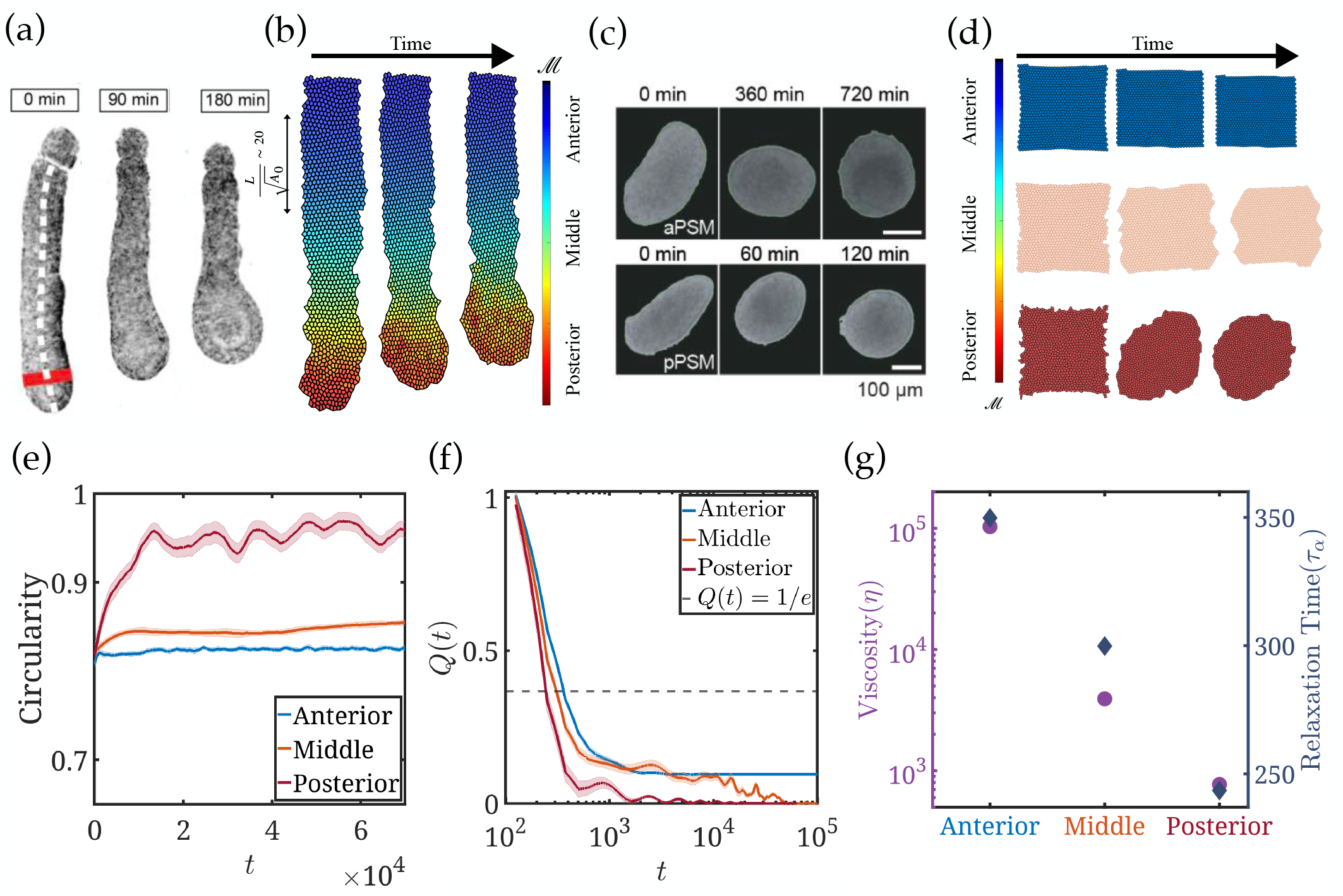
Effect of motility in presomitic mesoderm (PSM) rheology and relaxation. (a) Change in the shape of the PSM explant over time. Cell motility gradient drives pear-like shape formation by the explant [16] (Experimental image is taken with permission from [16]). (b) A numerical simulation of the vertex model with a gradient in random cell motility in the form of exponential decay [11] from the anterior to posterior axis has been introduced. The model qualitatively reproduces the experimental observation shown in (a). (c) Examples of rounding of anterior and posterior explants from PSM (image taken with permission from [16]). Anterior and posterior part shows significant distinction in rounding timescales, suggesting variation in viscosity along the A-P axis. (d) Numerical simulation of the vertex model of rounding of tissue from three different regions, anterior (blue), middle (green), and posterior (red) of the model PSM tissue. A higher cell motility results in faster rounding. (e) Time evolution of circularity of the tissue with three different motility values (as in (d)). (f) Time evolution of overlap function *Q*(*t*) (see Equation S8). It shows the rate of radial movement of a cell from its initial position. The dotted line marks when *Q*(*t*) drops below 1*/e* of its initial value. (g) Dependence of viscosity *η*, calculated from the Green-Kubo relation (see Materials and Methods for definition), and *α* relaxation Time *τα* on motility. The anterior region shows a high value of viscous timescales, and it drops linearly towards the posterior direction.

In our simulations, spreading arises from motility-driven junctional rearrangements, especially T1 transitions, where cells exchange neighbors. These rearrangements allow local structural reorganization. In regions with higher motility, such as the posterior PSM, cells more frequently overcome the energetic barriers associated with junctional remodeling, resulting in a more fluid-like behavior (Fig. S3). This inhomogeneous enhancement in tissue fluidity promotes inhomogeneous tissue deformation that causes the reproduction of experimentally observed morphology. Our results thus demonstrate active fluctuation driven rearrangements are sufficient to drive large-scale tissue remodeling in the absence of proliferation.

### B. Motility gradient leads to rheological heterogeneity along PSM

Experimental studies have suggested that variations in tissue viscosity along the A-P axis play a key role in shaping the morphogenesis of the PSM [2, 16]. To measure this rheological heterogeneity in our simulations, we quantified spatial variations in emergent tissue viscosity using two different approaches: the Green-Kubo formalism (see S.I., Sec. S2 C 2) and the *overlap function* approach (S.I. Sec. S2 B 1). The Green-Kubo formalism shows that the anterior region of the PSM exhibits significantly higher viscosity compared to the posterior, with viscosity decreasing along the AP axis (Fig. 1(g)). These results are in strong qualitative agreement with experimental measurements reported in Ref.[16]. Further analysis of cell dynamics across three distinct regions reveals that the *overlap function Q*(*t*), which measures the radial displacement of cells from their initial positions, exhibits three quantitatively distinct temporal decay profiles (Figure 1(f)). From the decay of *Q*(*t*), we estimate the relaxation time *τ*_*α*_ [see Materials and Methods for definition], which provides an estimate of the timescale over which tissue rearranges. We find that *τ*_*α*_ is shortest in the posterior region and longest in the anterior region [Fig. 1(g)], confirming that motility-driven fluidization is most prominent toward the posterior PSM. This gradient in viscous relaxation timescales reinforces the conclusion that tissue fluidity is spatially regulated and correlates with the observed morphological dynamics.

## IV. CELL MOTILITY CONTROLS TISSUE VISCOELASTICITY

To further dissect tissue rheology, we subjected the tissue to controlled shear deformations under varying levels of cellular motility. This approach allowed us to quantify fundamental rheological properties such as stress relaxation dynamics and the frequency-dependent storage and loss moduli, thereby establishing a direct link between local active fluctuations and the emergent viscoelastic behavior at the tissue scale.

### A. Stress Relaxation Test Reveals Motility Driven Residual Stress Retention

The anterior–posterior variation in tissue viscosity suggests that different regions of the PSM may respond differently to mechanical loading. To investigate how this rheological heterogeneity affects stress dissipation, we performed stress relaxation tests on tissues with varying motility levels. In each simulation, we applied a step shear by deforming the tissue affinely and fixing the outermost cell layers to maintain the applied strain. We then tracked the temporal evolution of shear stress in the bulk region, away from the boundary. By comparing tissues with low, medium, and high motility representing anterior, middle, and posterior PSM, we examined how active cellular dynamics influence the tissue’s ability to dissipate stress.

In all cases, we observed a characteristic biphasic relaxation behavior. At short to intermediate timescales, stress rapidly decayed (Fig. 2(a)). The slope of this early time relaxation phase was found to depend strongly on the cell motility: tissues with higher motility exhibited a steeper decay, indicating faster stress dissipation(Fig. 2(b)).

**FIG. 2.**
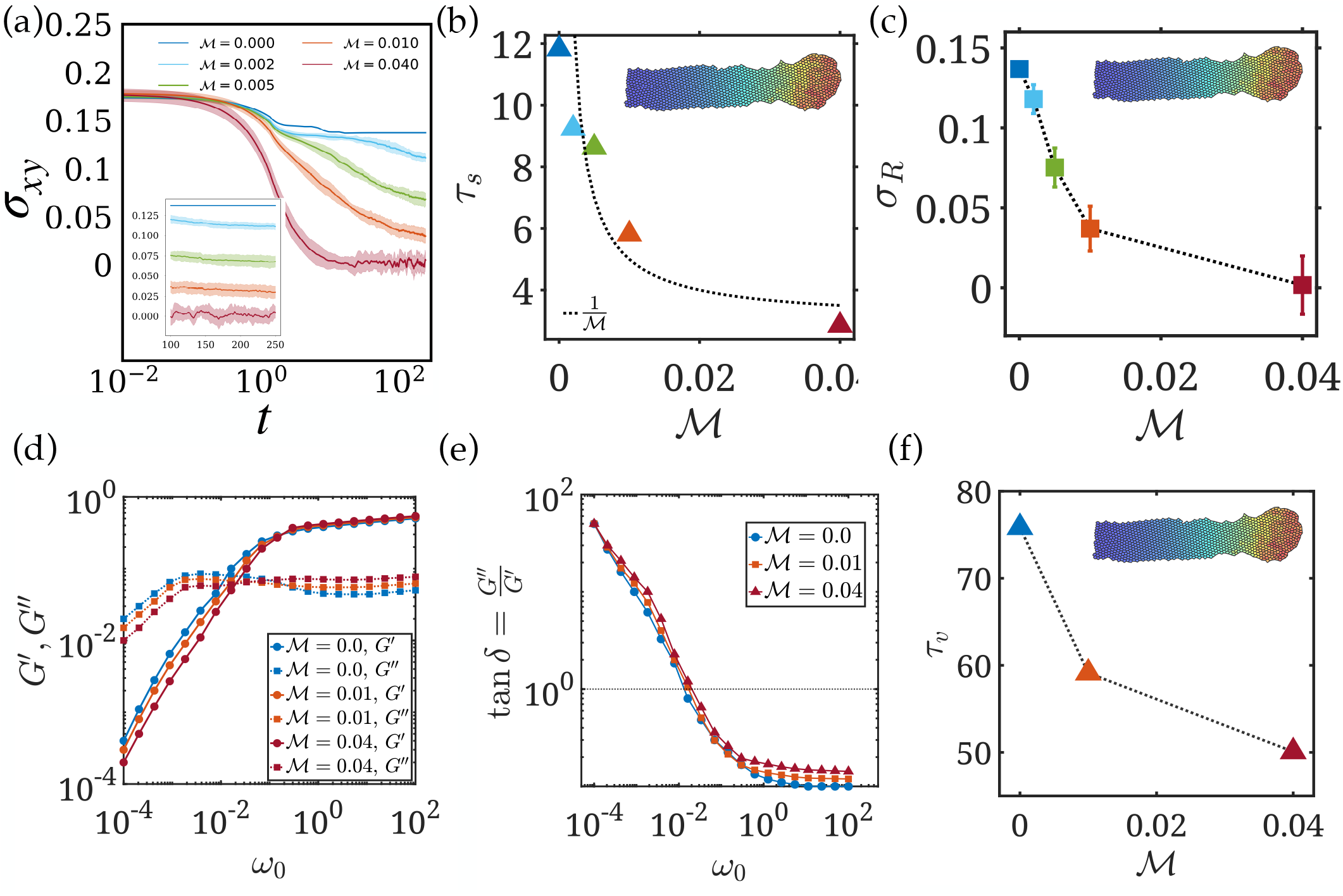
Tissue rheology under standard mechanical protocols. (a) Stress relaxation response for tissues with different motility levels. The decay of shear stress *σ*_*xy*_ (*t*) is shown following a step shear deformation. Higher motility leads to faster stress relaxation and lower residual stress. *Inset:* Long-time behavior of *σ*_*xy*_ reveals a nonzero plateau, indicating residual stress retention. (b) Relaxation timescale *τ*_s_ decreases with increasing motility, indicating enhanced fluidization. (c) Residual stress as a function of motility. More motile tissues retain less stress over time. (d) Frequency-dependent storage modulus *G*^*′*^ and loss modulus *G*^*′′*^. At low frequencies, viscous behavior dominates (*G*^*′′*^ *> G*^*′*^); at higher frequencies, the tissue responds more elastically. (e) The loss tangent tan δ = *G*^*′′*^*/G*^*′*^ as a function of driving frequency *ω*_0_, illustrating the transition from viscous to elastic dominance. (f) Intrinsic timescale of the tissue(*τ*_*v*_ ), defined by the crossover frequency where *G*^*′*^ = *G*^*′′*^, shifts to lower values with increasing motility, reflecting faster stress relaxation dynamics in more active tissues.

At longer timescales, the stress curves plateaued, revealing a non-zero residual stress in the tissue [Fig. 2(a), *Inset* ]. This behavior indicates solid-like features where cells cannot fully rearrange to relax all internal stresses. The magnitude of this residual stress was highest in the low-motility (anterior) regime and dropped as we went towards higher motility zones (posterior) [Fig. 2(c)]. This motility-dependent decline in residual stress qualitatively agrees with experimental observations in embryonic PSM where the posterior part shows lower residual stress than the anterior PSM (e.g. [2, 3]).

These stress relaxation test results suggest that motility not only governs the rate of stress dissipation but also tunes the mechanical state of the tissue from a more elastic, stress-retaining regime to a more fluidized, actively relaxing state. These findings help explain how developing tissues adjust their mechanical behavior during morphogenesis by tuning how actively their cells move and rearrange.

### B. Oscillatory Shear Reveals Cell Motility modulates Viscoelastic Crossover Frequency

To probe the viscoelastic properties of the tissue, we subjected the model tissue to low-amplitude oscillatory shear across a wide range of frequencies. This approach enabled us to extract the frequency-dependent storage modulus (*G*^*′*^), representing the elastic (energy-storing) response, and the loss modulus (*G*^*′′*^), representing the viscous (energy-dissipating) response. Together, these moduli provide a quantitative measure of the solid or fluid-like behavior of the tissue at different timescales of mechanical deformation.

At low oscillation frequencies, where the period of deformation exceeds the intrinsic stress relaxation time of the tissue, we observed that *G*^*′′*^ *> G*^*′*^. This indicates a viscosity-dominated response, in which the tissue can fully relax the imposed stress before the next deformation cycle begins, making the tissue behave more like a fluid. This corresponds to situations where cells have sufficient time to have enough junctional rearrangements (T1 transitions), contributing to tissue flow (Fig. 2(d)).

In contrast, at high frequencies, the oscillation timescale becomes shorter than the tissue’s internal relaxation time, leading to an increase in elastic behavior of the tissue (*G*^*′*^ *> G*^*′′*^, Fig. 2 (d)). Here, the tissue behaves more like a solid, storing deformation energy rather than dissipating it. The quick change of the shearing cycle does not give enough time for cellular junctional rearrangements or internal stress relaxation, causing the tissue to respond as an elastic material (Fig. 2(d)).

This fluid-to-solid transition is further quantified by the loss tangent tan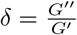, which decreases with frequency [Fig.2(e)]. The characteristic crossover frequency, defined by the condition *G*^*′′*^ = *G*^*′*^, marks the intrinsic relaxation timescale of the tissue. Importantly, we find that this crossover shifts to higher frequencies as cellular motility increases [Fig.2(f)], indicating that active cell movements facilitate more rapid stress relaxation.

This result highlights a key role for active cellular processes in modulating the viscoelastic landscape of the tissue, tuning its mechanical state between more fluid-like or solid-like regimes depending on the frequency of applied stress.

## V. TEMPORALLY PULSATED SPATIAL PERTURBATION GENERATES PERMANENT DEFORMATION IN VISCOELASTIC TISSUE

Having established the motility-dependent viscoelastic nature of the tissue, we next use the extracted viscoelastic timescales to predict tissue morphology under controlled external perturbations. Such analytical formulations offer a powerful means to uncover the fundamental principles underlying complex tissue dynamics. By abstracting the system into a continuum viscoelastic framework, we can extract scaling relations, isolate key parameter dependencies, and obtain predictions that remain broadly generalizable across contexts.

### A. Analytical Treatment Shows Motility Driven Viscoelasticity and Perturbation Length Scale Controls Permanent Tissue Deformation

Motivated by the presence of multiple relaxation timescales in the tissue, we model the synthetic vertex tissue as a two-dimensional viscoelastic medium using a linear viscoelastic model with two timescales[17]. This representation comprises a dashpot with viscosity *η*_1_ in series with a Kelvin–Voigt element, itself consisting of a spring with modulus *E* in parallel with a dashpot of viscosity *η*_2_ (Fig. 3(a) (I), (II)). The constitutive stress– strain relation for this system is given by

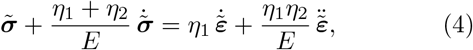

where 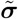 and 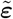 denote the stress and strain tensors, respectively. We then subject this material to an external perturbation that is both spatially patterned and temporally pulsed. Specifically, the forcing has a sinusoidal spatial profile of wavelength *k*_0_ and is applied in periodic on–off cycles, with duration T_on_ and T_off_ respectively (Fig. 3(a) III):

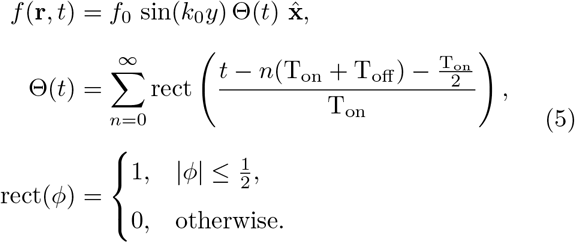

**FIG. 3.**
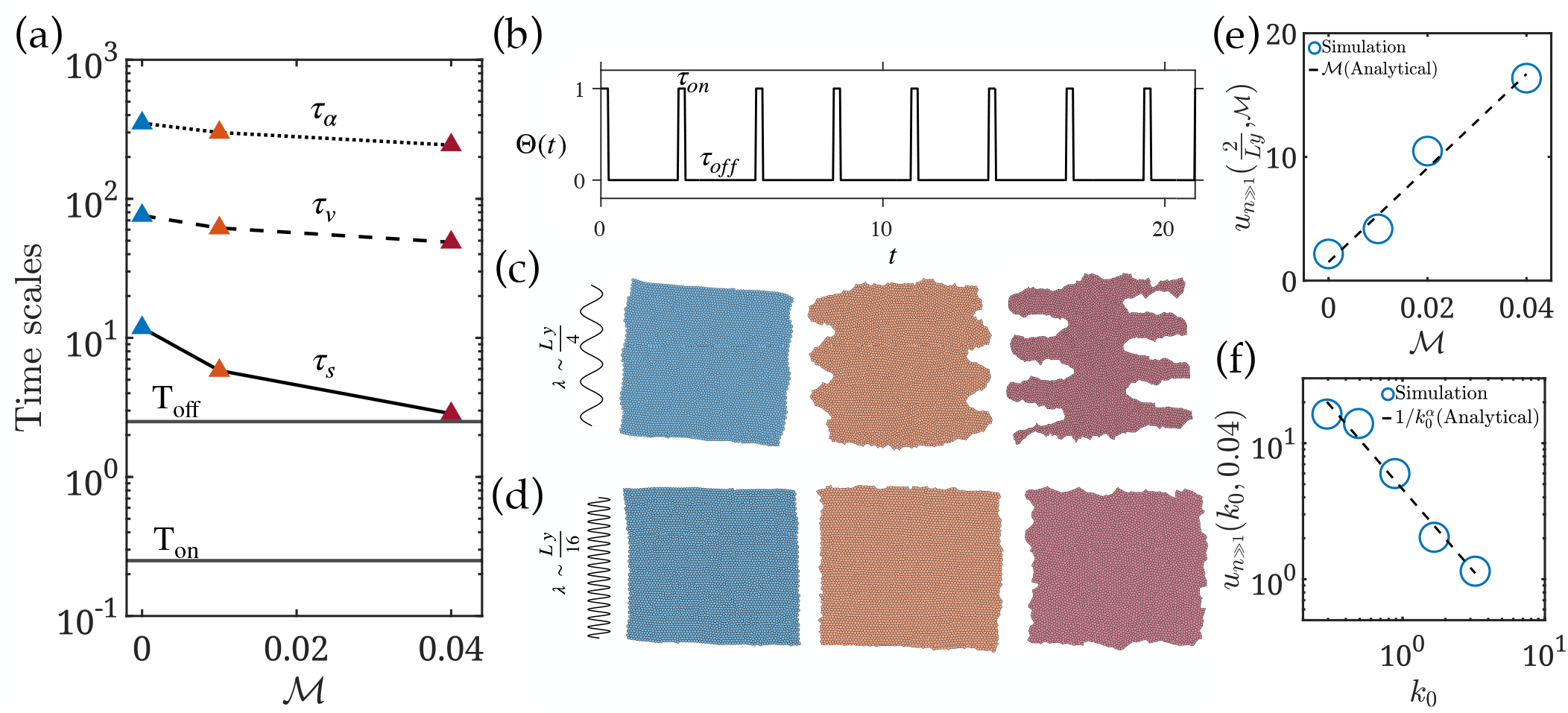
Tissue morphology under temporally pulsatile, spatially sinusoidal perturbations. (a) Schematic of the analytical framework. (I) Synthetic tissue used from the vertex model. (II) Continuum representation of the tissue as a 2D viscoelastic material, modeled as a dashpot (viscosity *η*_1_) in series with a Kelvin–Voigt element (dashpot with viscosity *η*_2_ in parallel with a spring of elasticity *E*). (III) Generic time profile of the external forcing: the force remains on for a duration T_on_ and off for a period T_off_ . (b) Comparison of characteristic timescales extracted from structural relaxation dynamics (*τ*_*α*_) and standard rheological protocols (*τ*_*s*_ and *τ*_*v*_ ) along the anterior–posterior (A–P) axis of the PSM. (c) Morphological outcomes of the vertex-model tissue under pulsatile forcing. (I) Low motility with long-wavelength perturbations ( ∼ *L*_*y*_*/*4, where *L*_*y*_ is tissue length along *y*) produces negligible morphological adaptation, but bulk rotation arises from force asymmetry. (II) Intermediate motility with the same wavelength induces moderate adaptation. (III) High motility yields pronounced morphological adaptation. (IV) Shorter-wavelength perturbations ( *L*_*y*_*/*16) fail to elicit significant adaptation or rotation. (V, VI) No morphological adaptation is observed. (d) Analytical predictions versus simulations of long-time tissue morphology in Fourier space, 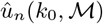, as a function of motility ℳ and wavenumber *k*_0_. Here *n* is the number of on-off cycles. *n* ≫ 1 is chosen. (I) Predicted deformation increases linearly with motility, in quantitative agreement with simulations. (II) Theory predicts a power-law decay of deformation with increasing wavenumber, matching the simulation findings.

**FIG. 4.**
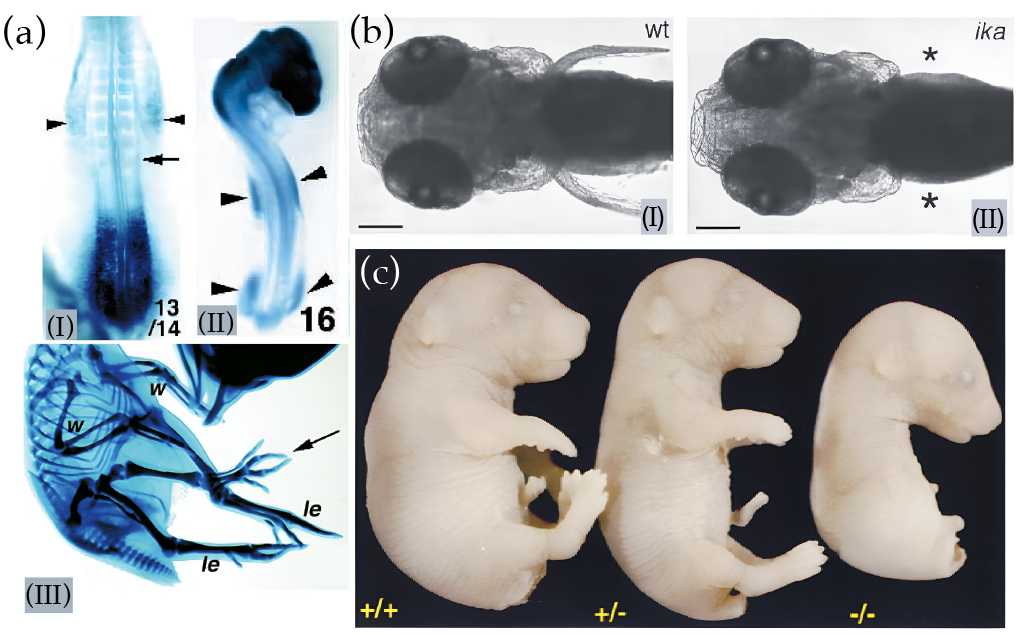
Effect of FGF signaling on vertebrate limb formation. (a) Chick embryo (adapted from Ohuchi et al. [18]): (I) Weak *Fgf10* expression in the prospective forelimb mesoderm (arrowheads) is seen around 13 Hamburger and Hamilton (HH) stage. (II) *Fgf10* expression in the head region and prospective limb mesoderm (arrowheads) increases at around the 16 HH stage. Suggesting *Fgf10* plays a key role in limb bud formation. (III) Induction of an ectopic leg-like limb (arrow) following implantation of FGF10-expressing cells in the interlimb region. (b) Zebrafish embryo (adapted from Fischer et al. [19]): (I) Wild-type larva at 3 dpf with pectoral fins protruding from the flanks. (II) *ika* mutant (Fgf24-deficient) larva lacking pectoral fins (asterisks). (c) Mouse embryo (adapted from Min et al.[20]): lateral views of *Fgf10* ^+*/*+^, *Fgf10* ^+*/*−^, and *Fgf10* ^−*/*−^ embryos, showing complete limb absence in the knockout.

We found the deformation at the end of one cycle (*t* = T_on_ +T_off_) is given in terms of the two timescales 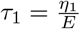 and 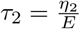 as

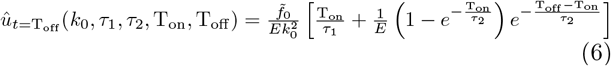

(See S.I. for detailed calculations.)

From this, we can extract the dependence of the deformation after *n* cycles, with one timescale *τ*_1_ and the perturbation wavelength *k*_0_ as,

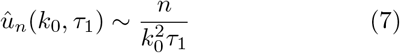

To directly connect the analytical deformation with experimentally accessible parameters such as motility, we first identify *τ*_1_ with the stress relaxation time, *τ*_*s*_. From Fig. 2(b), we established that *τ*_*s*_ scales inversely with motility, ℳ, i.e. *τ*_*s*_ ∼ 1*/* ℳ. Substituting this relation into the analytical expression yields a compact scaling form for the steady-state deformation amplitude:

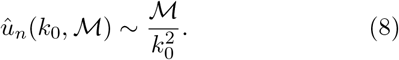

It is important to note that the analytical deformation, calculated at the end of a single T_off_ interval, represents the elementary contribution per cycle. Over multiple cycles, this deformation accumulates to set the long-term behavior; however, the underlying scaling laws with ℳ and *k*_0_ remain unaffected.

This expression reveals several key features. For any finite T_on_, the material exhibits a nonzero residual deformation at the end of each actuation cycle. Such residual deformations accumulate over successive cycles, potentially leading to lasting morphological transformations. In addition, the deformation amplitude scales linearly with the inverse of the fast relaxation timescale *τ*_1_. Finally, Relation 8 emphasizes the spatial dependence of the response: in the limit *k*_0_→ ∞ (corresponding to short-wavelength perturbations), the deformation amplitude vanishes. Together, these results predict that the timescale of cellular relaxation and the wavelength of external forcing jointly determine the efficiency of morphogenetic remodeling.

### B. Active Vertex Model Tissue Validates Analytical Prediction

To test these analytical predictions, we applied to our synthetic tissue a perturbation similar to that described in Eq. 5. Each cycle consisted of an active phase of duration *τ*_on_ followed by a passive phase of duration *τ*_off_, chosen such that *τ*_on_ *< τ*_*v*_≪ *τ*_*α*_ and *τ*_off_ ≲ *τ*_s_ (Fig. 3(b)). These parameters were selected as an extreme case relative to the analytical predictions to ensure that the simulated dynamics span the most separated timescales. This timing ensured that each active input was delivered within the tissue’s elastic response window, before complete stress relaxation. The forcing amplitude was kept below the intrinsic length scale of T1 transitions, thereby preventing the perturbation from directly inducing topological rearrangements. This setup allows us to isolate how short-lived external inputs applied in an elastic response window interact with slower internal relaxation processes to drive long-term tissue organization. Despite the short-lived nature of individual perturbations, our simulations show that tissues can undergo significant morphological changes over time, depending on both cell motility and the spatial scale of the perturbation.

For long-wavelength perturbations (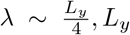is the length of the tissue along y direction), tissue behavior shows strong dependence on motility. In low motility regimes (ℳ ∼ 0.00), the tissue stress relaxation timescale (*τ*_s_) is much longer than the relaxation period of the applied perturbation (*τ*_off_) (Fig. 3(b)). Consequently, stress fails to dissipate neither via internal relaxation nor via junctional rearrangements. Instead, the asymmetric and cyclic nature of the sinusoidal forcing drives a coherent bulk rotation of the tissue, and no spatially structured patterns emerge. (Fig. 3(c)I). Intermediate motility ( ℳ= 0.01) allows some level of rearrangement but is insufficient to fully align with the imposed field, leading to partial morphological imprinting( Fig. 3(c) II). Only in high motility tissues (ℳ = 0.04), where *τ*_s_ ∼ T_off_ (Fig. 3(b)) junctional remodeling is efficient, and cells align locally with the imposed deformation(Fig.3(c)III) creating a strong morphogenetic pattern.

When pulsatile perturbations are applied at short wavelengths (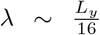, roughly four cell diameters), the tissue fails to exhibit any long-term morphological adaptation across all motility levels. Unlike longwavelength inputs, which impose global deformations, short-wavelength perturbations generate localized edge-length variations arising from the rapid spatial alternation of applied forces. These sharp local distortions increase the frequency of T1 transitions, allowing the tissue to rapidly relax back to its original configuration.

In the low-motility regime, one might expect slow stress relaxation to favor shape retention, as in the longwavelength case. However, here the imposed short-wavelength fluctuations directly drive local junctional rearrangements, overriding the slow viscoelastic response and preventing any stable morphological adaptation (Fig. 3(c)(IV)). For intermediate motility, junctional rearrangements occur even more rapidly, further accelerating local relaxation and again erasing any imposed pattern (Fig.3(c)(V)). At the highest motility levels, cells actively remodel their junctions, and the combination of intrinsic motility with edge-length fluctuations leads to transient small-scale boundary undulations. Yet, unlike the long-wavelength perturbations that generate coherent and lasting morphological changes, these fluctuations are quickly dissipated and do not translate into stable adaptations (Fig.3(c)(VI)).

Taken together, these results highlight a strong length-scale dependence in tissue mechanics. Long-wavelength perturbations couple to global viscoelastic modes, enabling robust, system-level adaptation, whereas short-wavelength perturbations are funneled into local junctional rearrangements and dissipated, leaving no lasting imprint.

We further performed a quantitative comparison between analytical predictions and simulation results. Specifically, we computed the long-term deformation amplitude in Fourier space, *u*_*n*_(*k*_0_, ℳ) (see S.I. Section S3 for details), and compared it against the theoretical scaling. The simulations show that the deformation amplitude increases linearly with motility ℳ, in agreement with the analytical prediction (Fig. 3(d)(I)). Also, both theory and simulations reveal a clear power-law dependence on perturbation length scale. While the theoretical prediction gives an exponent *α* = 2, the simulations yield a slightly smaller value, *α* ≈ 1.34 (Fig. 3(d)(II)), yet still capture the same underlying scaling behavior.

## VI. DISCUSSION

Our work provides a systematic framework for understanding how cell motility modulates the viscoelastic response of embryonic tissue such as PSM. Using an extended vertex model incorporating active forces, we characterized mechanical behavior under classical rheological protocols such as stress relaxation and oscillatory shear, as well as under spatiotemporally pulsatile perturbations. These studies reveal how motility modulates viscoelastic timescales, which can be used to understand pattern formation in embryonic tissue. We find that motility significantly accelerates stress relaxation, shifting tissue behavior from solid-like to fluid-like by enabling faster junctional rearrangements. In low-motility regimes, residual stress persists over long timescales, leading to elastic storage and global torque under repeated perturbation. In contrast, high-motility tissues reorganize locally, allowing for stress dissipation and morphological adaptation. These findings are consistent with in vivo observations in vertebrate presomitic mesoderm, where posterior regions exhibit fluid-like behavior due to elevated motility, while anterior regions behave more elastically [2, 9].

A minimal analytical framework captures the essential principles of tissue response under pulsatile forcing. Within a linear viscoelastic description [17], the model predicts that long-time deformation decays with increasing perturbation wavenumber (*k*_0_), reflecting the material’s intrinsic inability to retain high-frequency (cellular-scale) spatial features. This scale-dependent attenuation implies that tissues naturally act as mechanical filters, suppressing fine cellular-scale deformations while preserving longer-wavelength, sub-tissue level patterns.

To test these predictions, we investigated how transient, spatially structured perturbations—motivated by morphogenetic signals—affect long-term tissue morphology in the vertex model. Even when perturbations were applied within the elastic regime (T_on_ *< τ*_*v*_), tissues accumulated deformation across cycles. The resulting stress led to either alignment or global deformation, depending on the strength of motility and the spatial correlation length of the imposed cue. In agreement with the analytical predictions, sub-tissue–scale perturbations produced persistent morphological patterns in highly motile tissues, whereas cellular-scale cues were effectively dissipated or mechanically filtered out.

Taken together, our results reveal that the spatial correlation of the imposed cue and the level of cellular motility jointly determine the effective bandwidth over which tissues retain and respond to biochemical or mechanical inputs. The intrinsic viscoelasticity of the tissue thus governs how temporal and spatial features of external stimuli are integrated during morphogenesis. Notably, this framework establishes a mechanistic link between molecular signaling and large-scale mechanics: pathways such as FGF can modulate cell motility to tune viscoelastic responses, thereby shaping collective cell dynamics and morphogenetic outcomes.

FGF signaling has been proven to be a key controller of vertebrate limb development. In chick embryos, *Fgf10* expression emerges early in the prospective fore-limb mesoderm, initially weak around stage 13 (Hamburger and Hamilton (HH) 13) and becoming stronger by stage 16 (HH16), coinciding with the onset of limb bud outgrowth [18]. Functional perturbation experiments demonstrate causality: implantation of FGF10-expressing cells in the interlimb region induces ectopic, leg-like structures, underscoring FGF10’s sufficiency in initiating bud formation [18]. This role is conserved in zebrafish through Fgf24, where wild-type larvae develop prominent pectoral fins. At the same time, the *ika* mutant (lacking Fgf24) completely fails to initiate fin outgrowth [19], highlighting the requirement of FGF signaling for epithelial and mesenchymal motility. Similarly, in mouse embryos wild-type (*Fgf10* ^+*/*+^) and heterozygotes (*Fgf10* ^+*/*−^) develop normal limbs, homozygous knockouts (*Fgf10* ^−*/*−^) show complete limb absence [20]. Our results suggest that this FGF-mediated limb formation across species is governed by a localised increase in motility in otherwise static regions (e.g., lateral plate), which modulates the tissue’s viscoelastic properties locally, enabling directed outgrowth and limb formation.

Moreover, during embryogenesis, tissues are exposed to pulsatile mechanical cues spanning a wide range of spatial and temporal scales. Internally, many morphogenetic processes are driven by periodic contractile activity, for example, actomyosin pulses that drive apical constrictions [21] and the pulsed forces that underlie dorsal closure [22], as well as oscillatory gene expression that governs boundary formation during somitogenesis [23].

Tissues also experience rhythmic external forces. In early vertebrate embryos, the developing heart begins rhythmic contractions even before circulation, sending mechanical pulses across millimeter scales that can influence adjacent tissues, including the PSM [24–26]. Similarly, cilia-driven nodal flows generate oscillatory shear forces that not only direct left–right symmetry breaking but may also mechanically stimulate neighboring mesodermal regions [27]. Together, these internally generated and externally applied rhythmic inputs shape local cell behaviors and coordinate tissue-scale morphogenetic events such as axis elongation, boundary formation, and lumen expansion.

Also, the pulsatile force analysis developed here broadly applies beyond developmental morphogenesis. Many epithelial tissues in physiological settings experience rhythmic or cyclic mechanical loading. For example, the alveolar epithelium in the lungs undergoes continuous stretching and relaxation during normal breathing and mechanical ventilation, where the frequency and amplitude of deformation vary with respiration patterns [28– 30]. The intestinal epithelium is rhythmically deformed by peristaltic contractions that generate traveling waves of compression and extension along the gut [31]. Cardiac tissues are subjected to high-frequency cyclic strain during heartbeat cycles, with the heart epithelium enduring over 2.5 billion contraction and expansion events over an average human lifetime [32]. Other examples include bladder epithelial stretch during filling and voiding cycles, and uterine epithelium undergoing deformation during contractions.

In contexts such as mechanical ventilation, where the alveolar epithelium experiences cyclic stretching, deformation buildup over a long period may contribute to ventilator-induced lung injury. Conversely, tuning the frequency and amplitude of mechanical input to match tissue relaxation dynamics or modulating tissue fluidity through physiological or pharmacological interventions may reduce stress accumulation and preserve integrity. Similarly, our model predicts that tissues subject to rhythmic deformation selectively accumulate or dissipate stress depending on the match between input parameters and motility-driven remodeling capacity. This mechanistic framework thus links cyclic input characteristics with emergent morphological and stress responses, offering insight into how tissues can be tuned for robustness under repetitive mechanical loading.

This framework provides a versatile platform for investigating time-dependent pattern formation and stress encoding in active viscoelastic media. It can be readily extended to incorporate features such as anisotropic tension generation, active nematic alignment, and mechanochemical feedback, thereby enabling the exploration of more complex viscoelastic behaviors relevant to morphogenesis, organoid mechanics, and the design of programmable synthetic tissues.

## VII. ACKNOWLEDGEMENTS

These simulations were performed on the Param Seva supercomputers through the National Supercomputing Mission and the IITH Kanad clusters. S.I. acknowledges the Prime Minister Research Fellowship (PMRF, ID-2002732) for financial support through a research fellowship. M.S.R. acknowledges SERB(India) for financial support. A. G. acknowledges SERB-DST (India) Projects MTR/2022/000232, CRG/2023/007056-G, DST (India) grant no. DST/NSM/R&D HPC Applications/2021/05 and grant no. SR/FST/PSI-215/2016, and IITH for Seed Grant No. IITH/2020/09 for financial support.

