## Supplemental Information for "Motility-Driven Viscoelastic Control of Tissue Morphology in Presomitic Mesoderm"

### Motility driven viscoelasticity of embryonic tissue (Supplimentary Material)

#### S1. MATERIALS AND METHODS

##### A. Vertex Model

The energy functional and force for vertex model is given by,

$$U_c = \lambda(A_c - A_0)^2 + \beta P_c^2 + \gamma P_c \tag{S1}$$

$$\tag{S2}$$

The total force due to this energy functional on a vertex is given as:

$$\mathbf{F} = \begin{bmatrix} F_x \\ F_y \end{bmatrix} = -\nabla_v(U_c) = -2\lambda(A_c - A_0)\begin{bmatrix} \frac{\partial Ac}{\partial x_v} \\ \frac{\partial Ac}{\partial y_v} \end{bmatrix} -2\beta P_c \begin{bmatrix} \frac{\partial P_c}{\partial x_v} \\ \frac{\partial P_c}{\partial y_v} \end{bmatrix} - \gamma \begin{bmatrix} \frac{\partial P_c}{\partial x_v} \\ \frac{\partial P_c}{\partial y_v} \end{bmatrix}. \tag{S3}$$

The area of a polygon is given by:

$$A_c = \frac{1}{2}\sum_{v=1}^{N}(x_v y_{v+1} - y_v x_{v+1}),$$

where $v+1$ is the next vertex index, and the indices are cyclic (i.e., $x_{N+1} = x_1$).

The gradient of $A_c$ with respect to $x_v$ is:

$$\frac{\partial A_c}{\partial x_v} = \frac{1}{2}(y_{v+1} - y_{v-1}),$$

where $v+1$ and $v-1$ refer to the next and previous vertices, respectively.

Similarly, the gradient of $A_c$ with respect to $y_v$ is:

$$\frac{\partial A_c}{\partial y_v} = \frac{1}{2}(x_{v-1} - x_{v+1}).$$

The perimeter of a polygon is:

$$P_c = \sum_{v=1}^{N_v}\sqrt{(x_{v+1} - x_v)^2 + (y_{v+1} - y_v)^2}.$$

The gradient of $P_c$ with respect to $x_v$ involves contributions from both adjacent edges ($v-1 \to v$ and $v \to v+1$):

$$\frac{\partial P}{\partial x_v} = \frac{x_v - x_{v-1}}{\sqrt{(x_v - x_{v-1})^2 + (y_v - y_{v-1})^2}} + \frac{x_v - x_{v+1}}{\sqrt{(x_{v+1} - x_v)^2 + (y_{v+1} - y_v)^2}}$$

Similarly, the gradient of $P_c$ with respect to $y_v$ is:

$$\frac{\partial P_c}{\partial y_v} = \frac{y_v - y_{v-1}}{\sqrt{(x_v - x_{v-1})^2 + (y_v - y_{v-1})^2}} + \frac{y_v - y_{v+1}}{\sqrt{(x_{v+1} - x_v)^2 + (y_{v+1} - y_v)^2}}$$

Hence, the $x$- and $y$-gradients of $A_c$ and $P_c$ are:

$$\frac{\partial A_c}{\partial \mathbf{r}} = \begin{bmatrix} \frac{\partial A_c}{\partial x_v} \\ \frac{\partial A_c}{\partial y_v} \end{bmatrix} = \frac{1}{2}\begin{bmatrix} y_{v+1} - y_{v-1} \\ x_{v-1} - x_{v+1} \end{bmatrix} \tag{S4}$$

$$\frac{\partial P_c}{\partial \mathbf{r}} = \begin{bmatrix} \frac{\partial P_c}{\partial x_v} \\ \frac{\partial P_c}{\partial y_v} \end{bmatrix} = \begin{bmatrix} \frac{x_v - x_{v-1}}{\sqrt{\Delta x_{v-1}^2 + \Delta y_{v-1}^2}} + \frac{x_v - x_{v+1}}{\sqrt{\Delta x_{v+1}^2 + \Delta y_{v+1}^2}} \\ \frac{y_v - y_{v-1}}{\sqrt{\Delta x_{v-1}^2 + \Delta y_{v-1}^2}} + \frac{y_v - y_{v+1}}{\sqrt{\Delta x_{v+1}^2 + \Delta y_{v+1}^2}} \end{bmatrix} \tag{S5}$$

*

†

where $\Delta x_{v-1} = x_v - x_{v-1}$, $\Delta y_{v-1} = y_v - y_{v-1}$, and similarly for $\Delta x_{v+1}$ and $\Delta y_{v+1}$.

To make the cells motile we have added random white noise force $\xi_{c,v}(t)$ with $\langle \xi_{c,v}(t) \rangle = 0$ and $\langle \xi_{c,v}(t)\xi_{c',v'}(t') \rangle = 2\mathcal{M}\delta_{cc'}\delta_{vv}\delta(t-t'))$ to each vertex position($v$ of a cell $c$) that gives rice to random motion to the cells.

$$\eta \dot{\mathbf{r}}_c^v = -\nabla_v(U_c) + \boldsymbol{\xi}_{c,v}(t) \tag{S6}$$

This model incorporates cell junctional rearrangements (T1 transitions) and allows cells to detach from the tissue through T2 transitions or through multiple T1 transitions.

The parameters involved in equation S6 was non dimensionalized using the length scale $L \sim \sqrt{A_0}$ and time scale $\frac{1}{\sqrt{\lambda A_0}}$. Both the values of $\lambda$ and $A_0$ were kept 1 throughout all the simulations. The deterministic part of equation S6 was solved by implicit Euler time integration method and the stochastic part was solved by Weiner method with time step, $dt = 2.5 \times 10^{-3}$.

#### S2. ANALYSIS

##### A. Rheological Properties

###### 1. *Tissue circularity*

We have calulated tissue circularity by finding out the convex hull of the cell centres of the final structure. Then calculating the area ($A$) and perimeter($P$) of the given QHull the circularity is defined as,

$$\text{Circularity} = \frac{4\pi A}{P^2} \tag{S7}$$

##### B. Dynamic Properties

###### 1. *Overlap function*

The overlap function $Q(t)$ is defined by,

$$Q(t) = \left\langle \frac{1}{N_c} \sum_{c=1}^{N_c} W\left(a - |\mathbf{r}_c(t) - \mathbf{r}_c(0)|\right) \right\rangle \tag{S8}$$

Where , $W(x)$ is a Heaviside step function given by, $W(x<0) = 1$. It shows the rate of radial movement of a cell from its initial position. The relaxation time for a tissue is defined by the time it takes for $Q(t)$ to become $\frac{1}{e}^{th}$ of its initial value.

##### C. Rheological Properties

###### 1. *The stress tensor*

The stress tensor for an individual cell is calculated by,

$$\hat{\boldsymbol{\sigma}}_{\boldsymbol{c}} = -\Pi_c \hat{\mathbf{I}} + \frac{1}{2A_c} \sum_{e \in c} \boldsymbol{T}_e \otimes \boldsymbol{l}_e \tag{S9}$$

Where $\Pi_c = -\frac{\partial U_c}{\partial A_c}$ is the hydrostatic pressure and $\boldsymbol{T}_e = \frac{\partial U_c}{\partial \boldsymbol{l}_e}$ is the line tension/ shear stress term. This leads to the expression,

$$\hat{\boldsymbol{\sigma}}_{\boldsymbol{c}} = 2\lambda(A_c - A_0)\begin{pmatrix} 1 & 0 \\ 0 & 1 \end{pmatrix} + \frac{1}{2A_c}(2\beta P_c + \gamma) \sum_{v=1}^{N_v} \begin{pmatrix} \Delta x_{v+1}^2 & \Delta x_{v+1}\Delta y_{v+1} \\ \Delta y_{v+1}\Delta x_{v+1} & \Delta y_{v+1}^2 \end{pmatrix} \frac{1}{\sqrt{\Delta x_{v+1}^2 + \Delta y_{v+1}^2}} \tag{S10}$$

To calculate the stress tensor for the whole tissue, we do a weighted sum $\sigma = \sum_{c=1}^{N_c} \frac{A_c}{A_{total}} \sigma_c$.

###### 2. *Green-Kubo Viscosity*

To calculate viscosity of the system as an emergent phenomenon, we have used Green-Kubo relation,

$$\eta = \frac{A}{\mathcal{M}} \int_0^\infty < \sigma_{\alpha\beta}(0)\sigma_{\alpha\beta}(t) > \mathrm{dt} \tag{S11}$$

Here $A$ is the area of the system and $\mathcal{M}$ is the motility of the system. The Green-Kubo method estimates the viscosity from the autocorrelation of shear stress.

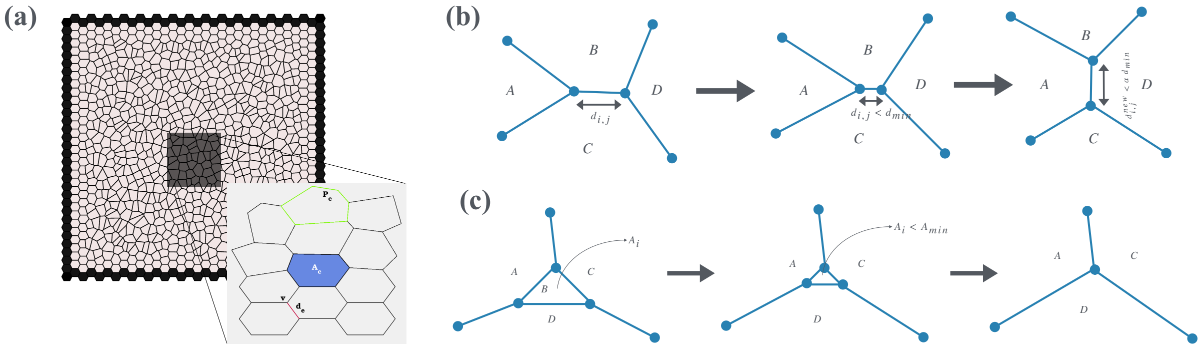


**Supplementary Figure S1:** (a) A representative epithelial tissue configuration obtained from vertex model simulations. (b) Schematic illustration of a T1 transition involving neighbor exchange. (c) Schematic of a T2 transition representing cell extrusion.

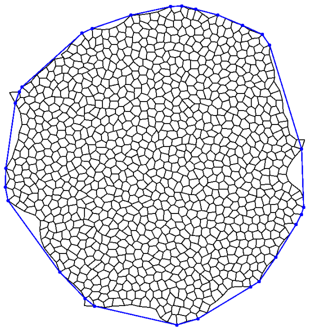

**Supplementary Figure S2:** Using convex hull to evaluate the circularity of tissue morphology.

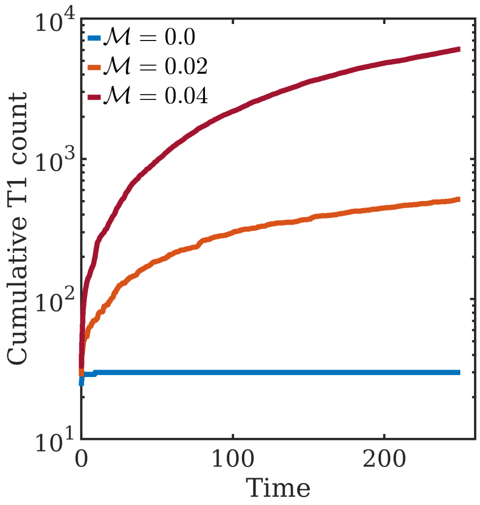


**Supplementary Figure S3:** Cumulative count of T1 transitions ocer time. Higher motility promotes T1 transitions.

##### *3. Stress relaxation experiment*

A stress relaxation experiment involves applying an external deformation to the system and observing how the internal stress dissipates over time. To achieve this in the simulation, we start form a random voronoi configuration and let the system relax just by it's own properties governed by the vertex Hamiltonian, and motility of the cells. Then we apply an affine shear strain $\gamma(t) = \begin{pmatrix} 1 & \epsilon \\ 0 & 1 \end{pmatrix}$ to the system and maintain the shear strain by keeping the boundary of the tissue fixed at a particular level of strain. We examine the relaxation of the shear stress of the bulk tissue, far away from the boundary.

##### *4. Oscillatory shear*

We have applied oscillatory shear strain to the system with varying frequency of the oscillation and measured the stress response of the system. From this we calculated Storage($G'$) and Loss modulus($G''$) given by the respectively in-phase and out-of-phase response of stress to the applied strain.

$$\begin{aligned} \epsilon(t) &= \epsilon_0 \sin(\omega_0 t) \\ \sigma(t) &= \sigma_0 \sin(\omega_0 t + \delta) \\ G' &= \frac{\sigma_0}{\epsilon_0} \cos(\delta) \\ G'' &= \frac{\sigma_0}{\epsilon_0} \sin(\delta) \end{aligned} \tag{S12}$$

*Calculation of $G'$ and $G''$ from $\sigma(t)$ :*
From equation S12, if we expand $\sigma(t)$ we get,

$$\begin{aligned} \sigma(t) &= \sigma_0 \left(\sin(\omega_0 t)\cos(\delta) + \cos(\omega_0 t)\sin(\delta)\right) \\ \sigma(t) &= \epsilon_0 G' \sin(\omega_0 t) + \epsilon_0 G'' \cos(\omega_0 t)) \end{aligned} \tag{S13}$$

Thus the we have to find out the Fourier coefficients corresponding to the input frequency $\omega_0$ from the Fourier series of $\sigma(t)$. One way is to do a *Fast Fourier Transform (FFT)* numerically and calculate the coefficients. But this requires large sampling data and can be errornous sometimes. To avoid that we have used an alternate calculation follows :

$$f(t) = a_0 + \sum_{n=1}^{\infty} a_n \cos(\omega_n t) + \sum_{n=1}^{\infty} b_n \sin(\omega_n t) \tag{S14}$$

Let's say $n = n_0$ corresponds to the input frequency. So we have to find the coefficients of $\cos(\omega_{n_0} t)$ and $\sin(\omega_{n_0} t)$ i.e $a_{n_0}$ and $b_{n_0}$.

Now, if we integrate equation S14 to a time $T$ or multiply it with $\cos(\omega_{n_0} t)$ or $\sin(\omega_{n_0} t)$ and then integrate, we get three equations involving $a_0$, $a_{n_0}$ and $b_{n_0}$ as three unknowns which we can solve as system of linear equations. These equations looks like,

$$\begin{bmatrix} \int_0^T f(t)\,dt \\ \int_0^T f(t)\cos(\omega_{n_0} t)\,dt \\ \int_0^T f(t)\sin(\omega_{n_0} t)\,dt \end{bmatrix} = \begin{bmatrix} \int_0^T 1\,dt & \int_0^T \cos(\omega_{n_0} t)\,dt & \int_0^T \sin(\omega_{n_0} t)\,dt \\ \int_0^T \cos(\omega_{n_0} t)\,dt & \int_0^T \cos^2(\omega_{n_0} t)\,dt & \int_0^T \sin(\omega_{n_0} t)\cos(\omega_{n_0} t)\,dt \\ \int_0^T \sin(\omega_{n_0} t)\,dt & \int_0^T \sin(\omega_{n_0} t)\cos(\omega_{n_0} t)\,dt & \int_0^T \sin^2(\omega_{n_0} t)\,dt \end{bmatrix} \begin{bmatrix} a_0 \\ a_{n_0} \\ b_{n_0} \end{bmatrix} \tag{S15}$$

In equation S15, the other terms where $n \neq n_0$ has been ignored as they would not contribute much. In equation S15, left hand side integrals are calculated using numerical integration scheme. Thus, we can calculate $a_0$, $a_{n_0}$ and $b_{n_0}$, and from that we can get $G' = \frac{b_{n_0}}{\epsilon_0}$ and $G'' = \frac{a_{n_0}}{\epsilon_0}$.

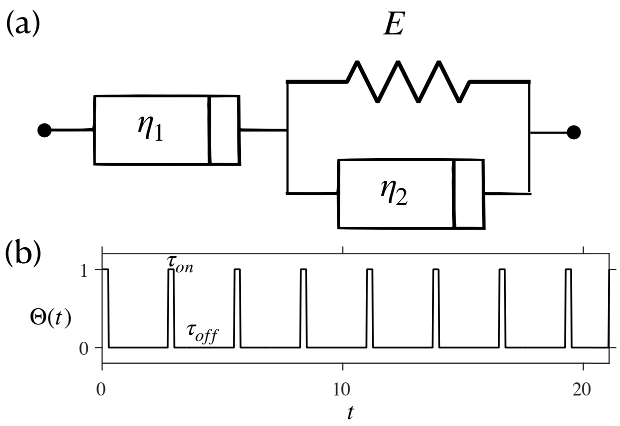


**Supplementary Figure S4:** (a) Standard Linear Fluid II model representing viscoelastic behavior, consisting of a dashpot with viscosity $\eta_1$ in series with a Kelvin–Voigt element (spring with modulus $E$ and dashpot with viscosity $\eta_2$). (b) Time profile of the applied mechanical perturbation.

#### S3. ANALYTICAL TREATMENT ( VISCOELASTIC MATERIAL UNDER PULSATILE SPATIAL PERTURBATION )

We investigated the response of a standard viscoelastic fluid subjected to spatially pulsatile perturbations, using the Standard Linear Fluid II model [1]. In this framework, a dashpot (viscosity $\eta_1$) is connected in series with a Kelvin–Voigt element, comprising a spring (modulus $E$) and a dashpot (viscosity $\eta_2$)(Fig S4,(a)). The resulting stress–strain relationship is given by:

$$\tilde{\boldsymbol{\sigma}} + \frac{\eta_1 + \eta_2}{E}\,\dot{\tilde{\boldsymbol{\sigma}}} = \eta_1\,\dot{\tilde{\boldsymbol{\varepsilon}}} + \frac{\eta_1 \eta_2}{E}\,\ddot{\tilde{\boldsymbol{\varepsilon}}} \tag{S16}$$

Where:

- $\tilde{\boldsymbol{\sigma}}$ is the stress tensor,

- $\tilde{\boldsymbol{\varepsilon}} = \frac{1}{2}\left(\nabla\mathbf{u} + \nabla\mathbf{u}^\top\right)$ is the linearized strain tensor,
- $\mathbf{u}(\mathbf{x}, t)$ is the displacement vector field,
- $E$ is the elastic modulus,
- $\eta_1$ and $\eta_2$ are the viscosity. (Fig. S4(a)).

This material is subjected to a pulsated perturbation (Fig. S4 (b)):

$$\Theta(t) = \sum_{n=0}^{\infty} \mathrm{rect}\left(\frac{t - n(\mathrm{T_{on}} + \mathrm{T_{off}}) - \frac{\mathrm{T_{on}}}{2}}{\mathrm{T_{on}}}\right) \tag{S17}$$

For $0 < t < \mathrm{T_{on}}$, the forcing is 1, and for $\mathrm{T_{on}} < t < \mathrm{T_{off}}$ the forcing is zero (Fig. S4(b)).

The force balance (quasistatic momentum conservation) is given by:

$$\nabla \cdot \tilde{\boldsymbol{\sigma}}(\mathbf{x}, t) + \mathbf{f}_{\mathrm{ext}}(\mathbf{x}, t) = 0, \tag{S18}$$

where $\mathbf{f}_{\mathrm{ext}}$ is the external body force per unit volume. Taking divergence of Equation S16, and defining,

$$\begin{aligned} a &= \frac{\eta_1\eta_2}{E} \\ b &= \eta_1 \\ d &= \frac{\eta_1 + \eta_2}{E}, \end{aligned} \tag{S19}$$

$$-\mathbf{f}_{\mathrm{ext}} - d\,\dot{\mathbf{f}}_{\mathrm{ext}} = b\,\partial_t(\nabla\cdot\tilde{\boldsymbol{\varepsilon}}) + a\,\partial_t^2(\nabla\cdot\tilde{\boldsymbol{\varepsilon}}) \tag{S20}$$

We consider the general 2D displacement field:

$$\mathbf{u}(x, y, t) = \begin{bmatrix} u_x(x, y, t) \\ u_y(x, y, t) \end{bmatrix}$$

From here,

$$\nabla\mathbf{u} = \begin{bmatrix} \partial_x u_x & \partial_y u_x \\ \partial_x u_y & \partial_y u_y \end{bmatrix}$$

And the strain tensor looks like:

$$\tilde{\boldsymbol{\varepsilon}} = \begin{bmatrix} \partial_x u_x & \frac{1}{2}(\partial_y u_x + \partial_x u_y) \\ \frac{1}{2}(\partial_y u_x + \partial_x u_y) & \partial_y u_y \end{bmatrix}$$

Hence,

$$-\begin{bmatrix} f_{\mathrm{ext},x} \\ f_{\mathrm{ext},y} \end{bmatrix} - d\begin{bmatrix} \dot{f}_{\mathrm{ext},x} \\ \dot{f}_{\mathrm{ext},y} \end{bmatrix} = \begin{bmatrix} b\,\partial_t\left(\partial_x^2 u_x + \frac{1}{2}(\partial_y^2 u_x + \partial_x\partial_y u_y)\right) + a\,\partial_t^2\left(\partial_x^2 u_x + \frac{1}{2}(\partial_y^2 u_x + \partial_x\partial_y u_y)\right) \\ b\,\partial_t\left(\partial_y^2 u_y + \frac{1}{2}(\partial_x^2 u_y + \partial_y\partial_x u_x)\right) + a\,\partial_t^2\left(\partial_y^2 u_y + \frac{1}{2}(\partial_x^2 u_y + \partial_y\partial_x u_x)\right) \end{bmatrix}$$

Since the forcing is in the $x$-direction and varies only with y, from the symmetry of the system, we make the ansatz for displacement field:

$$u_x = u(y, t) \quad u_y = v(y, t),$$

This implies:

$$\partial_x u_x = 0, \quad \partial_x^2 u_x = 0, \quad \partial_y\partial_x u_y = 0$$

The remaining non-zero derivatives are:

$$\partial_y^2 u_x = \partial_y^2 u(y, t), \quad \partial_t\partial_y^2 u_x = \partial_t\partial_y^2 u(y, t)$$

We know, $f_{\mathrm{ext},x} = f_x$ and $f_{\mathrm{ext},y} = 0$.

So, the force balance reads,

$$\begin{aligned} -f_x - d\,\partial_t f_x &= \frac{b}{2}\,\partial_t\partial_y^2 u(y, t) + \frac{a}{2}\,\partial_t^2\partial_y^2 u(y, t) \\ 0 &= b\ \partial_t\partial_y^2 v(y, t) + b\partial_t\partial_y^2 v(y, t) \end{aligned} \tag{S21}$$

We observe from Eqn S21 that the deformation in the y direction ($v(y, t)$) will decay with time.
To calculate the deformation in the x direction ($u(y, t)$), we need to solve Eqn S21. We start by taking a Fourier transform to eliminate the spatial derivatives and solve the ODE in time in Fourier space.

In Fourier space, the x-diraction force balance equation becomes (dropping the subscript x)

$$\begin{aligned} -\hat{f}(k, t) - d\,\partial_t\hat{f}(k, t) &= -\frac{k^2}{2}\left(b\,\partial_t\hat{u}(k, t) + a\,\partial_t^2\hat{u}(k, t)\right) \\ \implies \quad a\,\partial_t^2\hat{u}(k, t) + b\,\partial_t\hat{u}(k, t) &= \frac{2}{k^2}\left(\hat{f}(k, t) + d\,\partial_t\hat{f}(k, t)\right) \end{aligned} \tag{S22}$$

Now, we have $f(\mathbf{r}, t) = f_0\sin(k_0 y)\Theta(t)$. So, $\hat{f}(k, t) = f_0 \cdot \pi\frac{1}{i}\left[\delta(k + k_0) - \delta(k - k_0)\right]\Theta(t)$.

Hence, Eqn S22 reads,

$$a\,\partial_t^2\hat{u}(k, t) + b\,\partial_t\hat{u}(k, t) = \frac{f_o\pi}{ik^2}\left[\delta(k + k_0) - \delta(k - k_0)\right]\left[\Theta(t) + d\,\dot{\Theta}(t)\right] \tag{S23}$$

We observe that the forcing term on the right-hand side of Equation S23 is zero for any $|k| \neq k_0$. Since we are interested in the system's response to the external forcing, we restrict our attention to the mode $k = k_0$. Thus, we write Equation S22 specifically for the $k_0$ mode as:

$$a\,\partial_t^2 \hat{u}(k_0,t) + b\,\partial_t \hat{u}(k_0,t) = \frac{\pi f_0}{i k_0^2}\left(\Theta(t) + d\,\dot{\Theta}(t)\right) \quad \text{(S24)}$$

Now, we are going to focus on the deformation in one cycle ($t = \mathrm{T_{on}} + \mathrm{T_{off}}$), and also redefine the amplitude of forcing as complex amplitude $\tilde{f}_0 = \frac{\pi f_0}{i}$. So, within one cycle, the equation Eqn S24 becomes,

$$a\,\partial_t^2 \hat{u}(k_0,t) + b\,\partial_t \hat{u}(k_0,t) = \frac{\tilde{f}_0}{k_0^2}\left[\theta(t) + d\left(\delta(t) - \delta(t-\tau_{\rm on})\right)\right] \quad \text{(S25)}$$

Here, $\theta(t) = 1$ for $0 < t < \mathrm{T_{on}}$, and $\delta(t)$ is the dirac delta function. For notational simplicity we redefine, $c = \frac{\tilde{f}_0}{k_0^2}, d \equiv \frac{\tilde{f}_0}{k_0^2} d$. So, we have,

$$a\,\partial_t^2 \hat{u}(k_0,t) + b\,\partial_t \hat{u}(k_0,t) = c\;\theta(t) + d\;\left(\delta(t) - \delta(t-\tau_{\rm on})\right) \quad \text{(S26)}$$

The boundary conditions for solving Eqn S26 are,

$$\hat{u}(k_0,0) = 0, \quad \partial_t \hat{u}(k_0,0^+) = \frac{d}{a},$$
$$\hat{u}(k_0,\mathrm{T_{on}}^-) = \hat{u}(k_0,\mathrm{T_{on}}^+), \quad \partial_t \hat{u}(k_0,\mathrm{T_{on}}^+) - \partial_t \hat{u}(k_0,\mathrm{T_{on}}^-) = -\frac{d}{a} \quad \text{(S27)}$$

For $0 < t < \mathrm{T_{on}}$

$$\begin{aligned} \hat{u}_{\rm on}(k_0,t) &= C_1 + \frac{aC_2}{b} e^{-\frac{b}{a}t} + \frac{c}{b} t \\ \partial_t \hat{u}_{\rm on}(k_0,t) &= -C_2 e^{-\frac{b}{a}t} + \frac{c}{b} \end{aligned} \quad \text{(S28)}$$

And for $\mathrm{T_{on}} < t < \mathrm{T_{off}}$

$$\begin{aligned} \hat{u}_{\rm off}(k_0,t) &= C_3 + \frac{aC_4}{b} e^{-\frac{b}{a}(t-\mathrm{T_{on}})} \\ \partial_t \hat{u}_{\rm off}(k_0,t) &= -C_4 e^{-\frac{b}{a}(t-\mathrm{T_{on}})} \end{aligned} \quad \text{(S29)}$$

After applying the boundary conditions (Eqn S27), the coefficients $C_1, C_2, C_3$ and $C_4$ becomes,

$$\begin{aligned} C_2 &= \frac{c}{b} - \frac{d}{a} \\ C_1 &= -\frac{aC_2}{b} \\ C_4 &= C_2 e^{-\frac{b}{a}\mathrm{T_{on}}} - \frac{c}{b} + \frac{d}{a} \\ u_{\rm on}(\tau_{\rm on}) &= C_1 + \frac{aC_2}{b} e^{-\frac{b}{a}\mathrm{T_{on}}} + \frac{c\mathrm{T_{on}}}{b} \\ C_3 &= u_{\rm on}(\mathrm{T_{on}}) - \frac{aC_4}{b} \end{aligned} \quad \text{(S30)}$$

Finally, with all the coefficients known, we can find the deformation at $t = \mathrm{T_{off}}$,

$$\hat{u}_{t=\mathrm{T_{off}}}(k_0,a,b,C_3,C_4) = C_3 + \frac{a}{b} C_4 e^{-\frac{b}{a}(\mathrm{T_{off}}-\mathrm{T_{on}})} \quad \text{(S31)}$$

Since we are interested in the behavior of the final deformation with perturbation length scale and viscosity, after putting in the respective values of $C_1, C_2, C_3$ and $C_4$ in Eqn S31, we can get,

$$\hat{u}_{t=\mathrm{T_{off}}}(k_0,\tau_1,\tau_2,\mathrm{T_{on}},\mathrm{T_{off}}) = \frac{\tilde{f}_0}{E k_0^2}\left[\frac{\mathrm{T_{on}}E}{\eta_1} + \frac{1}{E}\left(1 - e^{-\frac{E}{\eta_2}\mathrm{T_{on}}}\right) e^{-\frac{E}{\eta_2}(\mathrm{T_{off}}-\mathrm{T_{on}})}\right] \quad \text{(S32)}$$

Identifying the two timescales $\tau_1 = \frac{\eta_1}{E}$ and $\tau_2 = \frac{\eta_2}{E}$, we rewrite the expression, as

$$\hat{u}_{t=\mathrm{T_{off}}}(k_0,\tau_1,\tau_2,\mathrm{T_{on}},\mathrm{T_{off}}) = \frac{\tilde{f}_0}{E k_0^2}\left[\frac{\mathrm{T_{on}}}{\tau_1} + \frac{1}{E}\left(1 - e^{-\frac{\mathrm{T_{on}}}{\tau_2}}\right) e^{-\frac{\mathrm{T_{off}}-\mathrm{T_{on}}}{\tau_2}}\right] \quad \text{(S33)}$$

From this, we can extract the dependence of the deformation with one timescale $\tau_1$ and the perturbation wavelength $k_0$ as,

$$\hat{u}_{t=\mathrm{T_{off}}}(k_0,\tau_1) \sim \frac{1}{k_0^2 \tau_1} \quad \text{(S34)}$$

Since, this residual deformation is going to accumulate in every cycle without affecting any scaling behavior. So we can write the total deformation at the end of cycle $n$,

$$\hat{u}_{n\gg 1}(k_0,\tau_1) \sim \frac{n}{k_0^2 \tau_1} \quad \text{(S35)}$$

To establish equivalence with our simulations, we identify the stress relaxation timescale $\tau_s$ from simulations with the viscous relaxation time $\tau_1$ in the Standard Fluid model. This identification is supported by the matched temporal decay of stress in both simulation data and analytical solutions, confirming that $\tau_s$ and $\tau_1$ characterize the same relaxation process. Based on this, the defomation becomes,

$$\hat{u}_{n\gg 1}(k_0, \tau_1) \sim \frac{1}{k_0^2 \tau_s} \tag{S36}$$

Note, we drop the $n$ as it is a constant.

Furthermore, since we have shown in the main text that motility $\mathcal{M}$ scales inversely with the stress relaxation time, i.e., $\mathcal{M} \sim 1/\tau_s$, we can re-express the deformation amplitude as:

$$\hat{u}_{n\gg 1}(k_0, \mathcal{M}) \sim \frac{\mathcal{M}}{k_0^2} \tag{S37}$$

This highlights how increasing motility enhances the tissue's mechanical response to spatial perturbations.

[1] P. Kelly, Mechanics Lecture Notes: An introduction to Solid Mechanics., Auckland: The University of Auckland (2025).
